# Production of diverse beauveriolide analogs in closely related fungi: a rare case of fungal chemodiversity

**DOI:** 10.1101/2020.07.01.183541

**Authors:** Ying Yin, Bo Chen, Shuangxiu Song, Bing Li, Xiuqing Yang, Chengshu Wang

## Abstract

Fungal chemodiversity is well known in part due to the production of diverse analogous compounds by a single biosynthetic gene cluster (BGCs). Usually, similar metabolites are produced by closely related fungal species. Here we report a rare case of the production of the cyclodepsipeptide beauveriolides (BVDs) in three insect pathogenic fungi. We found that the more closely-related fungi *Beauveria bassiana* and *B. brongniartii* produce structurally distinct analogs of BVDs whereas the rather divergently evolved species *B. brongniartii* and *Cordyceps militaris* produce structural analogs in a similar pattern. It was verified that a conserved BGC containing four genes is responsible for BVD biosynthesis in three fungi including a polyketide synthase (PKS) for the production of 3-hydroxy fatty acids (FAs) with chain length variations. In contrast to BVD production patterns, phylogenetic analysis of the BGC enzymes or enzyme domains largely resulted in the congruence relationship with fungal speciation. Feeding assays demonstrated that a FA with a chain length of eight carbon atoms was preferentially utilized whereas a FA with a chain longer than 10 carbon atoms could not be used as a substrate for BVD biosynthesis. We also found that addition of *D-*type amino acids could not enable *B. bassiana* to produce those analogs biosynthesized by other two fungi. Insect survival assays suggested that the contribution of BVD to fungal virulence might be associated with the susceptibility of insect species. The results of this study enrich the knowledge of fungal secondary metabolic diversity.

**IMPORTANCE:** Fungal chemotaxonomy is an approach to classify fungi based on fungal production of natural compounds especially the secondary metabolites. We found an atypical example that could question chemical classification of fungi in this study: the more closely-related entomopathogenic species *Beauveria bassiana* and *B. brongniartii* produce structurally different analogs of the cyclodepsipeptide beauveriolides whereas the rather divergent species *B. brongniartii* and *Cordyceps militaris* biosynthesize similar analogs under the same growth condition. The conserved BGC containing four genes is present in each species and responsible for beauveriolide production. In contrast to the compound formation profiles, the phylogenies of biosynthetic enzymes or enzymatic domains show associations with fungal speciation relationship. Dependent on insect species, production of beauveriolides may contribute to fungal virulence against insect. The findings in this study augment the diversity of fungal secondary metabolisms.

---

It is common that different analogs of certain secondary metabolites can be produced by a fungal species or different species of related or unrelated fungi (1, 2). For example, more than 30 structural analogs of cyclosporin A, the immunosuppressant drug, have been identified from the opportunistic insect pathogenic fungus *Tolypocladium inflatum* and other fungi (3). For insecticidal and phytotoxic destruxins, around 40 structural analogs with variations in bioactivities can be similarly produced by *Metarhizium* and other fungal species (4). With the elucidation of cyclodepsipeptide biosynthetic mechanisms, it is clear that the non-ribosomal peptide synthetase (NRPS) can uptake the alternate building blocks of amino acids or hydroxy acids for cyclodepsipeptide biosynthesis (5, 6). Taken together with the function of various tailoring enzymes, diverse cyclodepsipeptide analogs can therefore be produced by a single biosynthetic gene cluster (BGC) (2). It is still rare to find that the structurally-different analogs are produced by the conserved BGC in closely-related fungi.

The cyclodepsipeptide beauverolides were first identified in 1977 from the insect pathogenic fungus *Beauveria bassiana* (7). Different analogs were later identified either from *B. bassiana* or from its close relatives *B. tenella* (8), *Cordyceps militaris* (9), and *Paecilomyces* (*Isaria*) *fumosoroseus* (8, 10). However, the term beauveriolide instead of beauverolide was used for those analogs later identified in *C. militaris* or *Beauveria* sp. (9, 11). For simplicity, if not otherwise specified, the terms <u>b</u>eau<u>v</u>eroli<u>d</u>e and <u>b</u>eau<u>v</u>erioli<u>d</u>e are designated as BVD that are composed of a 3-hydroxy-4-methyl fatty acid [i.e. either 3-hydroxy-4-methyldecanoic acid (HMDA, 10 carbon atoms) or 3-hydroxy-4-methyloctanoic acid (HMOA, 8 carbon atoms)], two *L*-type and one *D*-type amino acid (AA) residues (Fig. 1A). Currently, there are 28 structurally different BVDs have been identified (Table 1). Given that alternate BVDs were isolated from the closely-related fungi, it is still unclear regarding the association of BVD productions with fungal speciation relationship.

**FIG 1.**
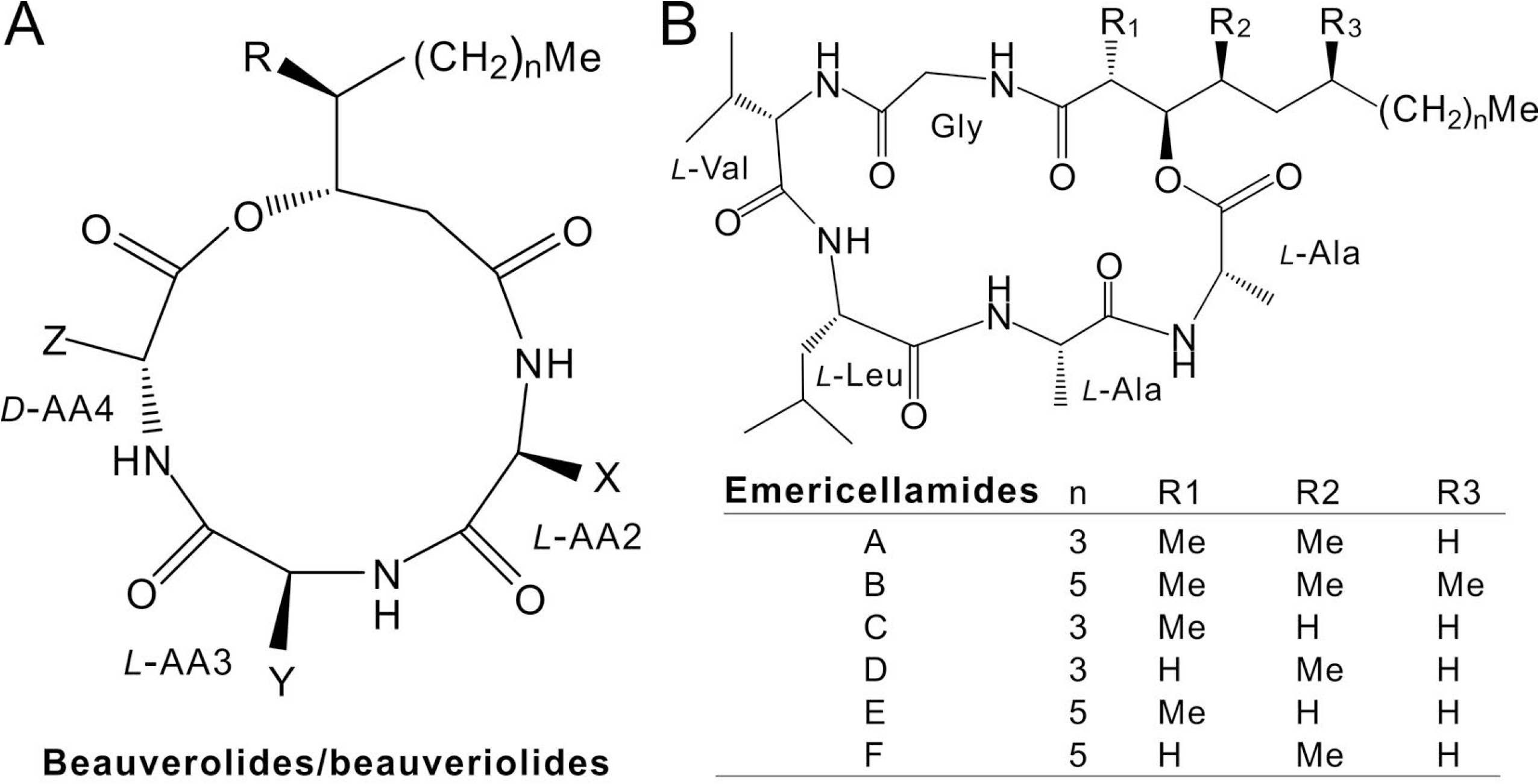
Structure comparison between BVDs and emericellamides. (A) Common structure of BVDs. The analogs with different amino acid residues *L-*AA2, *L*-AA3 and *D*-AA4 as well as the letters X, Y, Z, n and R can be referred in Table 1. (B) Chemical structures of emericellamides.

**Table 1.**
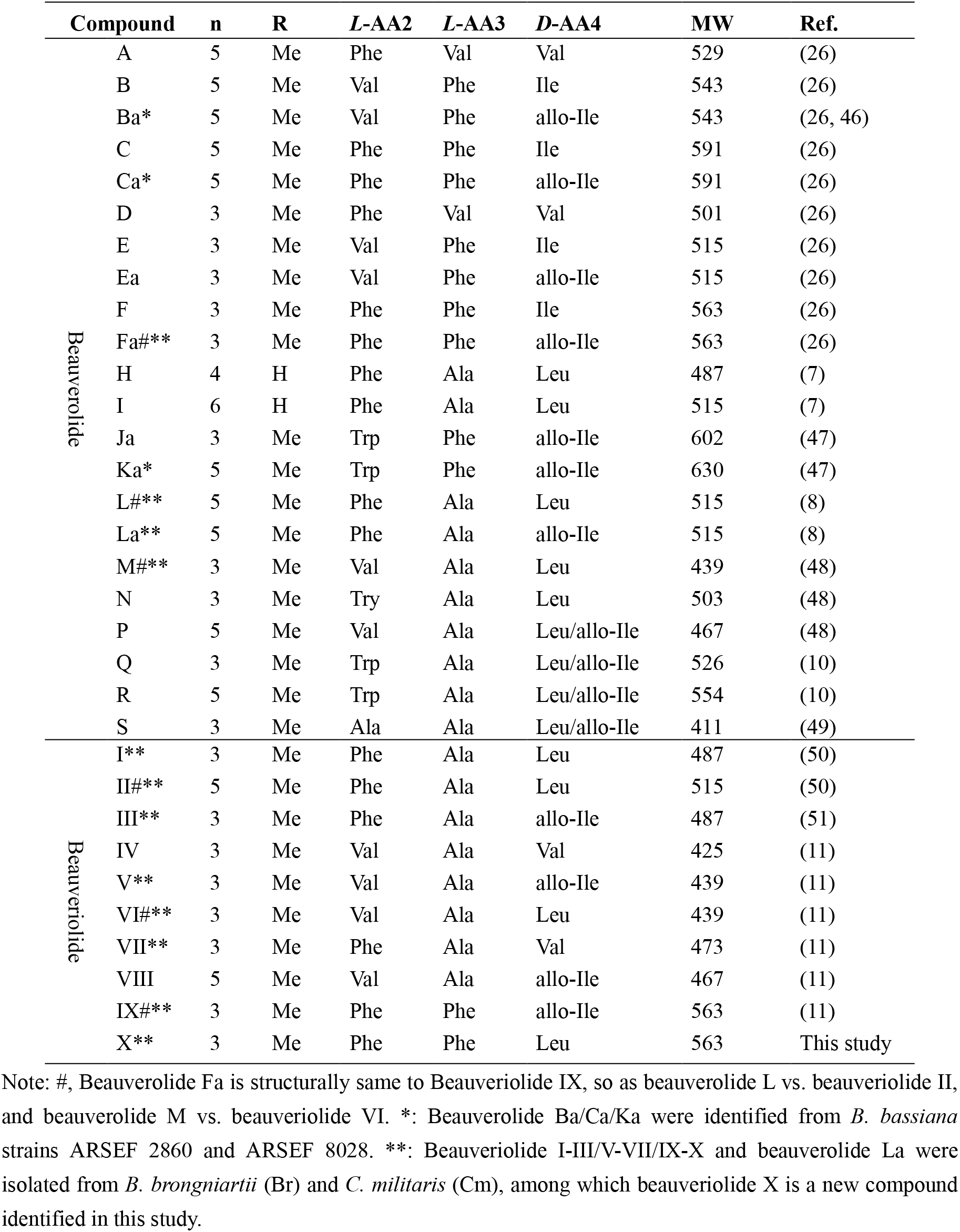
Composition of BVDs identified before and in this study.

Different biological and or medicinal activities have been reported for BVD analogs, including antiaging (12), beta-amyloid-lowering (13), and inhibition of acyl-CoA:cholesterol acyltransferase activity to block the synthesis of cholesteryl esters (14, 15). The analog of BVDs could be detected in insect hemolymph after *B. bassiana* infection (16). It has also been shown that injection of the wax moth (*Galleria mellonella*) larvae with beauverolide L could not kill insects in a dosage less than 30 micrograms per larvae but induce immune responses in insects (17). It is still unclear whether BVD production is required for full fungal virulence against insect hosts.

Chemical synthesis of BVD analogs has been successful (18). However, until this study, the genetic chemistry of BVD biosynthesis remains unclear. During our ongoing studies, a recent paper has reported the heterologous expression of the clustered genes from *C. militaris* to successfully detect the production of beauveriolides I and III in *Aspergillus nidulans* (9). We have obtained the genome information of BVD producing fungi (19–21). In this study, BVD production and gene deletions were conducted in three Cordycipitaceae species of insect pathogens including *B. bassiana*, *B. brongniartii* and *C. militaris*. It is interesting to find that the conserved BGC is responsible for the biosynthesis of different BVD analogs irrespective of fungal phylogenetic associations. Substrate feeding and insect bioassays have also been performed to postulate the substrate specificity of different enzymes or domains, and to determine the BVD contribution to fungal virulence against insects.

## RESULTS

### Prediction and structure analysis of the gene cluster

Considering that the cyclodepsipeptide BVDs contain a 3-hydoxy fatty acid (FA) as one of the building blocks like emericellamides produced by *A. nidulans* (Fig. 1B), the putative BGC for BVD biosynthesis was deduced to contain both a polyketide synthase (PKS) and a NRPS genes. Based on the reciprocal Blast analysis with the emericellamide biosynthetic genes *easA-easD* (22), similar to a previous analysis (9), a conserved BGC containing four genes is present in the genomes of three entomopathogenic fungi including *B. bassiana*, *B. brongniartii* and *C. militaris* (Table S1). The genes were termed *besA*-*besD* (for <u>be</u>auveriolide/<u>b</u>eauverolide <u>s</u>ynthesis) for those in *B. bassiana*, which is highly conserved to those in *B. brongniartii* and *C. militaris* but with gene order and orientation differences to the emericellamide BGC of *A. nidulans* (Fig. 2A). A reciprocal Blast analysis of the orthologous protein identities indicated that the enzymes of *B. bassiana* are mostly similar to those of *B. brongniartii* (84-95% at amino acid level) followed by those of *C. militaris* (69-81%) and *A. nidulans* (40-54%) (Table S1), indicating that the enzymes of two *Beauveria* species are more conserved to each other than those between *Beauveria* and *Cordyceps* fungi even all these species belong to the Cordycipitaceae fungi (23, 24).

**FIG 2.**
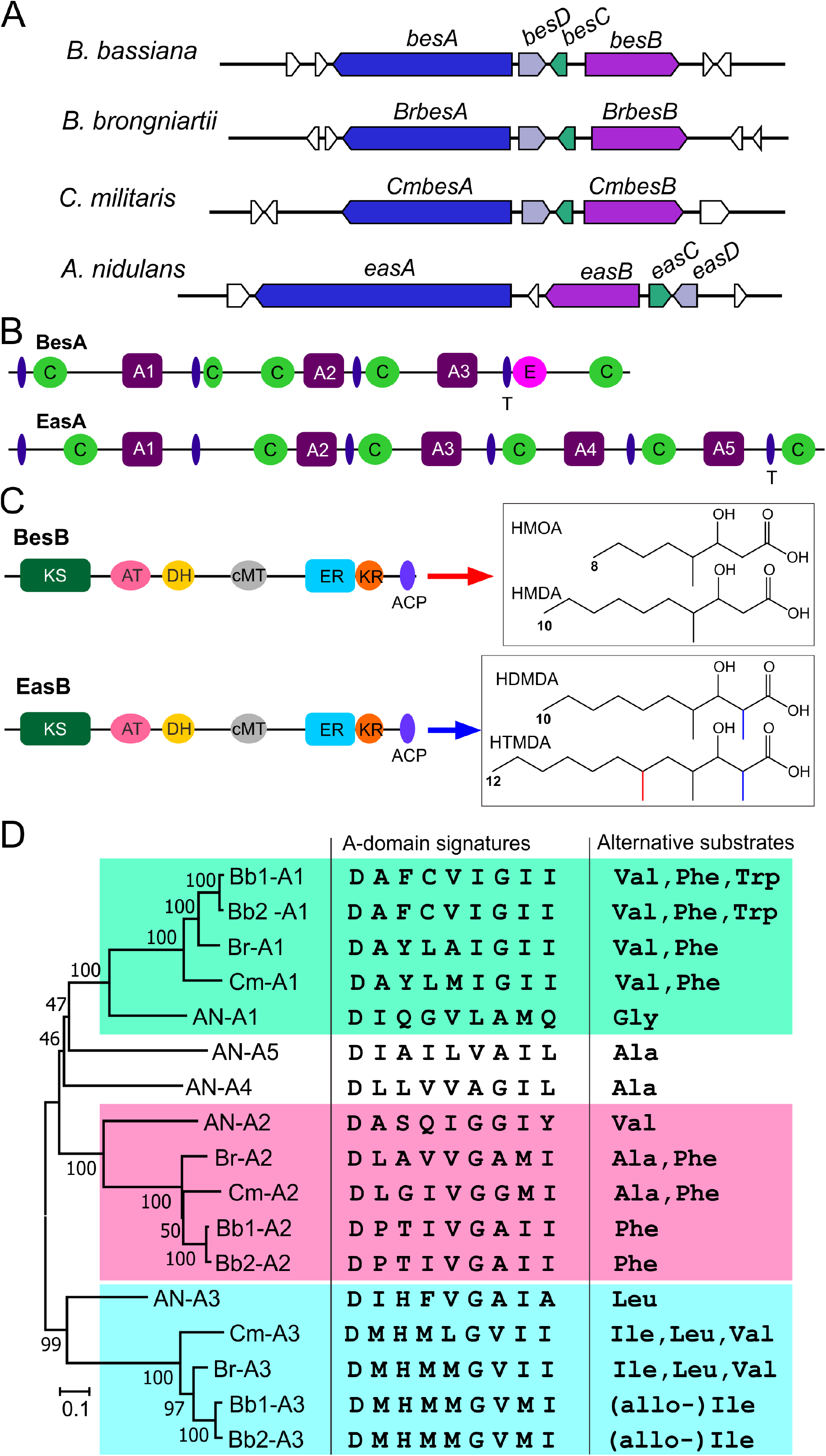
Conservation and phylogenetic analysis of gene cluster and core biosynthetic enzymes. (A) Conservative relationship of the gene clusters responsible for the biosynthesis of BVDs and emericellamides in different fungi. The genes labeled in the same color show orthologous relationships. (B) Schematic structure comparison between the NRPS enzymes BesA and EasA. Different domains are: C, condensation; A, adenlylation; E, epimerization; T, thiolation. (C) Schematic structure comparison between the PKS enzymes BesB and EasB. Different domains: KS, ketosynthase; AT, acyl transferase; DH, dehydratase; cMT, C-methyltransferase; ER, enoyl reductase; KR, ketoreductase; ACP, acyl-carrier-protein. Fatty acids are: HMDA, 3-hydroxy-4-methyldecanoic acid; HMOA, 3-hydroxy-4-methyloctanoic acid; HDMDA, 3-hydroxy-2,4-dimethyldecanoic acid; and HTMDA, 3-hydroxy-2,4,6-trimethyldodecanoic acid. (D) Phylogenetic, signature and alternative substrate analysis of the NRPS A-domains from different fungi. Fungal species or strains: Bb1, *B. bassiana* strain ARSEF 2860; Bb2, *B. bassiana* ARSEF 8028; Br, *B. brongniartii*; Cm, *C. militaris*; AN, *A. nidulans*.

Consistent with the structure difference between BVDs and emericellamides, the linear NRPS BesA is short of two modules when compared with EasA (Fig. 2B). In contrast, the PKS structures are similar between BesB and EasB (54% identity), however, the former might be responsible for the biosynthesis of HMOA and HMDA whereas EasB produces longer chain FAs of 3-hydroxy-2,4-dimethyldecanoic acid (HDMDA) and 3-hydroxy-2,4,6-trimethyldodecanoic acid (HTMDA) (Fig. 2C). Phylogenetic analysis of the adenlylation (A) domains retrieved from homologous NRPS enzymes suggest that the terminal A4- and A5-domain associated modules of EasA would have lost in BesA and its homologs in *B*. *brongniartii* and *C. militaris* (Fig. 2D). Except for the A2-domain, the A1- and A3-domain phylogenetic associations are following the fungal speciation trajectory, i.e., *A. nidulans* diverged first and then the Cordycipitaceae fungi in the order of *C. militaris*, *B. brongniartii* and *B. bassiana* (19). However, in terms of the A-domain signatures and alternative substrates deduced from the BVDs produced by respective fungal species (Table 1), *B. brongniartii* and *C. militaris* rather than *B. brongniartii* and *B. bassiana* might produce more similarly structured analogs.

### Verification of the BGC genes for BVD biosynthesis in different fungi

Having shown that the putative BGC is present in the genome of three closely-related fungal species, the strains of these three fungi were first incubated in a liquid culture and the mycelia were extracted with methanol for high performance liquid chromatography (HPLC) analysis. Intriguingly, consistent with A-domain signature analysis (Fig. 2D), the data revealed that the chromatographic profiles are similar to each other between *C. militaris* and *B. brongniartii*, which were different from the similar patterns produced by two strains of *B. bassiana* (Fig. 3A), i.e. the more closely-related *B. bassiana* and *B. brongniartii* produced divergent compounds.

**FIG 3.**
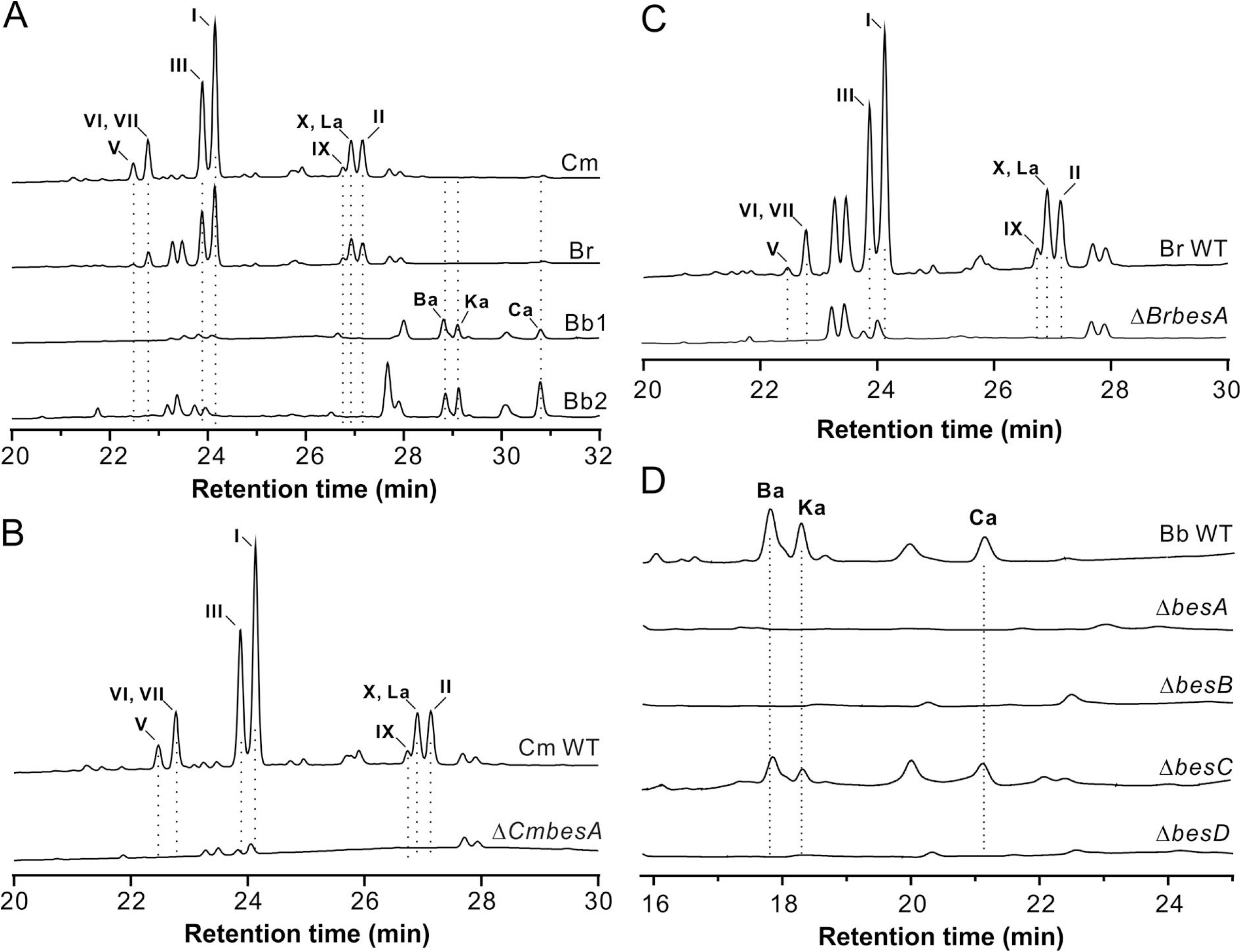
HPLC profiling of BVD production by the WT and null mutants of different fungi. (A) Production of BVD analogs by different fungal species. (B) Non-production of different BVDs by *C. militaris* after deletion of *CmbesA*. (C) Non-production of different BVDs by *B. brongniartii* after deletion of *BrbesA*. (D) HPLC profiling of BVD production after deletion of the clustered genes in *B. bassiana*.

After deletion of the core genes in each species, multiple peaks disappeared in the samples extracted from three fungi (Fig. 3B-3D). To determine whether these peaks are BVD analogs, the strains of *B. bassiana* and *B. brongniartii* were used for large-scale fermentations in Sabouraud dextrose broth (SDB) and metabolite purifications. The individually purified compounds were subject to nuclear magnetic resonance (NMR) analysis (Table S2-S13). As a result, 12 BVDs with alternate 3-hydroxy FAs (i.e., either HMDA or HMOA) and AA residues were identified in this study (Fig. S1). A new compound was identified after 2D NMR analysis (Fig. S2; Table S13), which is composed of HMOA, *L*-Phe, *L-*Phe and *D-*Leu with a structure different from 28 BVDs that have been identified from different fungi (Table 1). This compound is termed beauveriolide X that can only be detected from *C. militaris* and *B. brongniartii* when the fungi were grown in SDB. In terms of the structures, *B. bassiana* only produced three detectable HMDA-type BVDs Ba, Ka and Ca after examining two different strains. In contrast, *B. brongniartii* and *C. militaris* mostly biosynthesized HMOA-type BVDs (seven out of nine, with BVDs I and III being dominant) and two HMDA-type BVDs La and II (Fig. 3A; Table S14).

Contrary to *Beauveria* species (25), it is easy to induce sexual fruiting bodies for *C. militaris* (20). Thus, the fruiting bodies of *C. militaris* were induced both on silkmoth pupae and rice medium (Fig. S3A). In contrast to a previous report (9), metabolite extraction and HPLC analysis indicated that no detectable BVDs were accumulated in both types of fruiting-body samples (Fig. S3B).

### Putative biosynthetic pathway of BVDs

To further propose the biosynthetic pathway, the putative PKS gene *besB*, acyltransferanse *besC* and CoA-ligase *besD* were also individually deleted in *B. bassiana*. Similar to the null mutant of NRPS gene, Δ*besB* and Δ*besD* lost abilities to produce BVD Ba, Ka and Ca when compared with the wild type (WT) strain. However, relative to WT, a reduced amount of BVDs could still be produced by Δ*besC* (Fig. 3D). Thus, the biosynthetic pathways of BVDs could be suggested that BesB is responsible for the production of the 3-hydroxy FAs HMDA or HMOA by interactively catalyzing chain elongation using either three or four copies of manonyl-CoA. After the catalysis of CoA ligation by BesD, BesC may transfer CoA-HMDA or CoA-HMOA to the thiolation (T) domain of BesA for stepwise integration of alternate AAs to form different analog of BVDs (Fig. 4). It is noteworthy that based on all reported BVDs (Table 1), the upload of *D*-Ile by A3-domain to form BVD B, C, E or F in *B. bassiana* (26) has not been detected in this study after examining two different genotypic strains.

**FIG 4.**
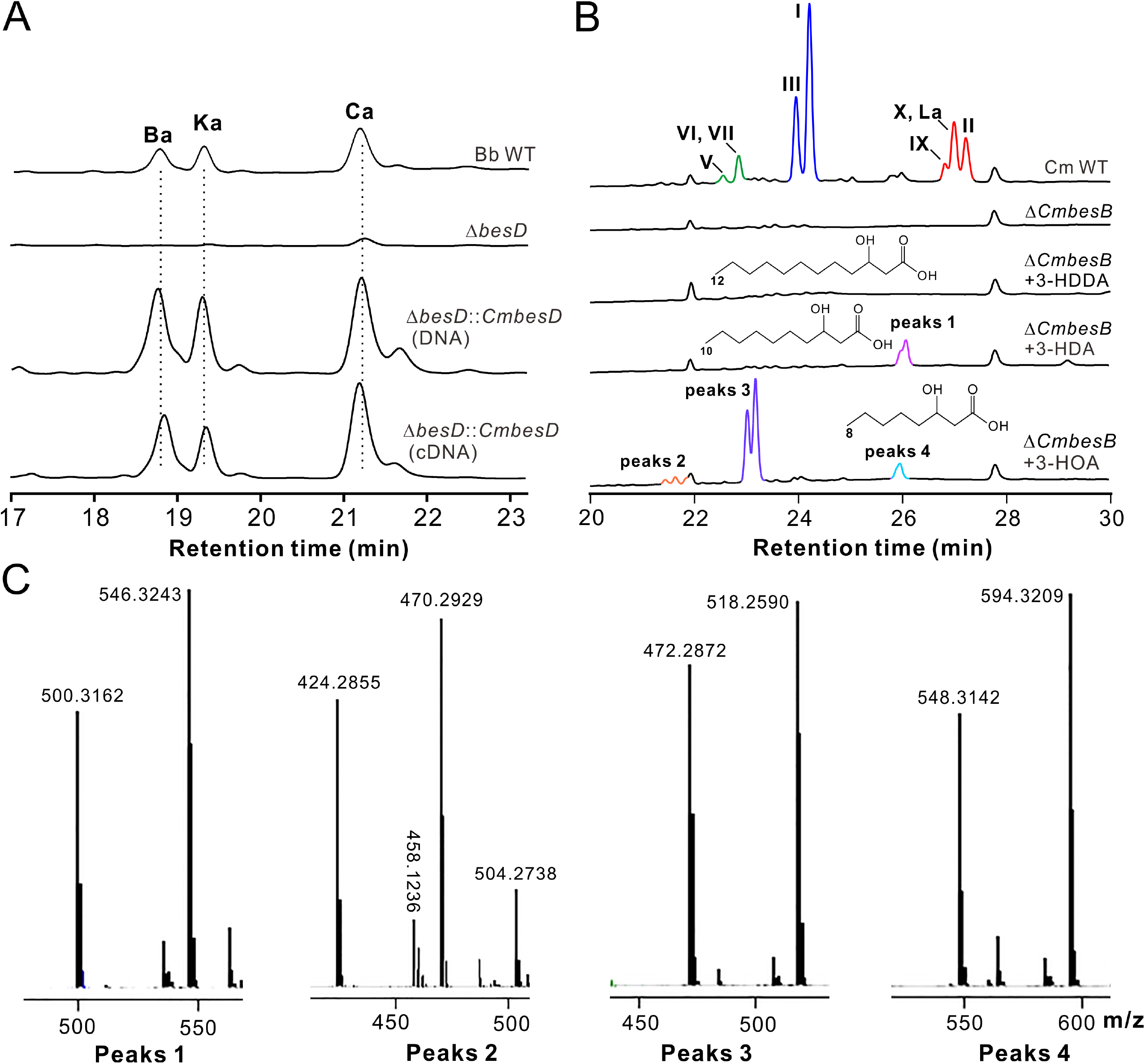
Schematic pathway for BVD biosynthesis. Different domains of the PKS enzyme are as shown in Fig. 2. Alternative substrates for the stepwise NRPS assembly line are framed for the production of BVDs detected in this study. The previsions of *D-*type AAs for module 4 are still unclear. The analogous structures with the letters X, Y, Z, n and R can be referred in Table 1.

### Substrate specificities for alternate 3-hydroxy FAs and *D*-type amino acids

As indicated above, *B. bassiana* produced different structure of BVDs from those biosynthesized by *C. militaris* and *B. brongniartii* (Fig. S1). We were curious about the alternative biosynthesis and integration of 3-hydroxy FAs, i.e., either HMDA or HMOA, for BVD productions in different fungi. The ketosynthase (KS), acyl transferase (AT), ketoreductase (KR) and enoyl reductase (ER) domain sequences of BesB and its homologs were retrieved for phylogenetic analysis. The tailoring enzyme BesC and BesD homologous sequences were also analyzed. It was found that, in contrast to the BVD production profiles (Fig. 3A), the phylogenetic relationships of both PKS domains and tailoring enzymes are all similarly associated with fungal speciation relationship, i.e., the domains and enzymes of *B. bassiana* and *B. brongniartii* are more closely related to each other than to those of *C. militaris* (Fig. S4).

We next performed gene-replacement test by complementation of Δ*besD* with its orthologous gene *CmbesD* from *C. militaris*. As a result, the gene rescue with either genomic DNA or cDNA of *CmbesD* could restore the ability of Δ*besD*, however, to only produce BVDs Ba, Ka and Ca like the WT strain of *B. bassiana* (Fig. 5A). We also tested the feeding assays of PKS gene mutants with the available 3-hydroxy FAs, i.e., 3-hydroxydodecanoic acid (3-HDDA, 12 carbon atoms), 3-Hydroxydecanoic acid (3-HDA, 10 carbon atoms) and 3-hydroxyoctanoic acid (3-HOA, 8 carbon atoms). The feeding of Δ*besB* did not detect any extra peak(s) when compared with the null mutant (Fig. S5A). However, detectable peaks could be obtained after feeding Δ*CmbesB* with 3-HAD and 3-HOA but not with the longer chain FA 3-HDDA. Intriguingly, more peaks were obtained after feeding with 3-HOA than with 3-HDA (Fig. 5B). The respective peaks were collected and subject to LC-mass spectrometry (MS) analysis. Based on the obtained mass spectra of both [M-H]^+^ and [M+COOH]^−^ for each peak (Fig. 5C), 3-HDA and 3-HOA might be incorporated into the NRPS assembly line to produce demethyl (dm) BVDs. For example, the feeding with 3-HDA might lead to the formation of putative dm-BVD L and or dm-BVD La (both with a molecular weight of 501) in Δ*CmbesB* of *C. militaris*. The feeding with 3-HOA could result in the production of at least seven dm-BVDs (Table S15) with the putative dm-BVDs I and III (both with a molecular weight of 473) being dominant (Fig. 5B and 5C), which is consistent with the preference of using HMOA for BVD productions in *C. militaris* (Table S14).

**FIG 5.**
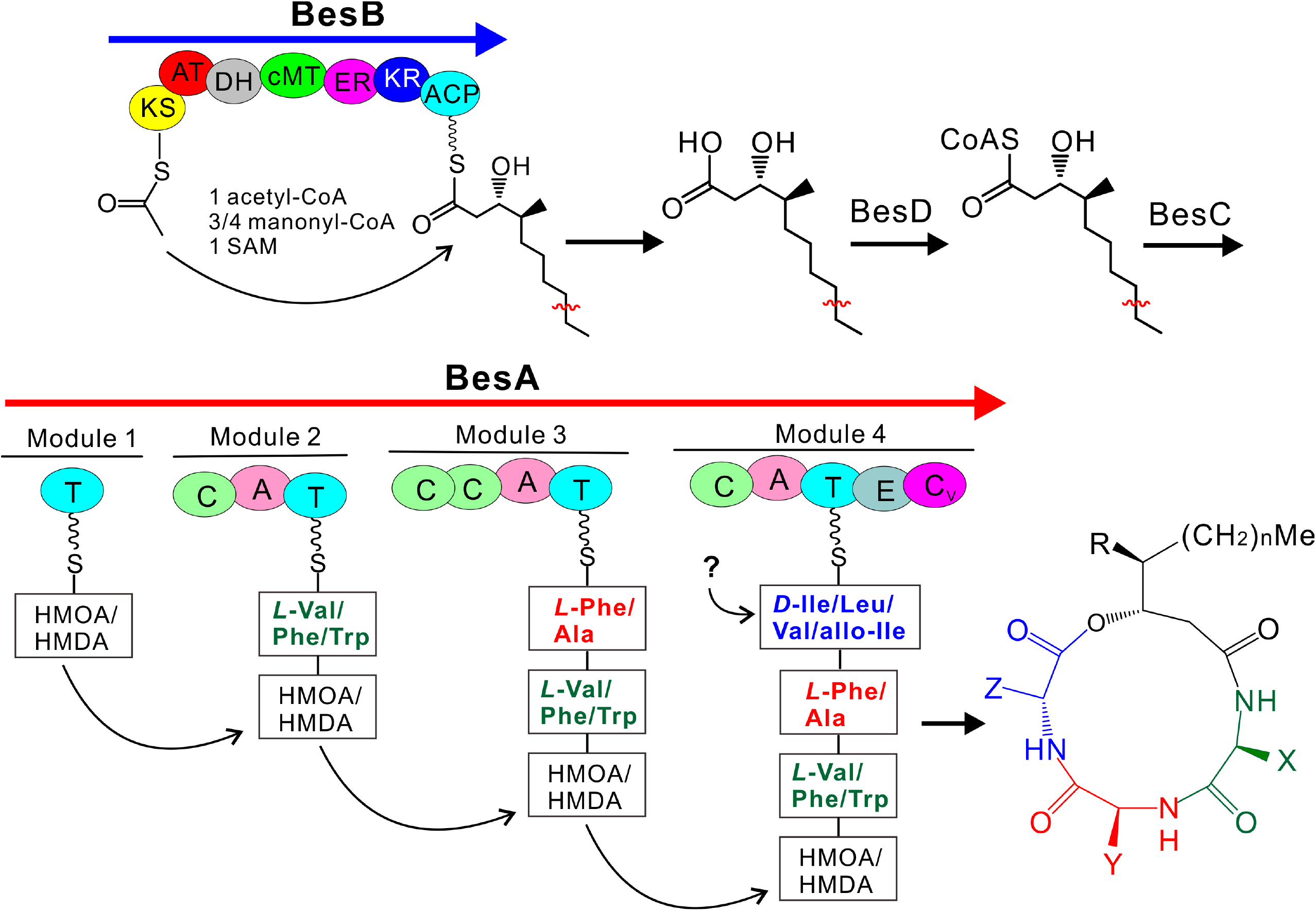
Substrate specificity tests. (A) HPLC profiling of compound production after complementation of Δ*besD* with the *CmbesD* gene. (B) HPLC profiling of metabolite production by Δ*CmbesB* after feeding with different dm-hydroxy FAs. Each FA was added at a final concentration of 100 μg/ml. FAs are: 3-HDDA, 3-hydroxydodecanoic acid; 3-HAD, 3-Hydroxydecanoic acid; 3-HOA, 3-hydroxyoctanoic acid. (C) Mass spectra (both [M-H]^+^ and [M+COOH]^−^) of the peaks detected after FA feedings of Δ*CmbesB* shown in panel B.

The BVDs produced by *B. bassiana* also differ from those of *B. brongniartii* and *C. militaris* in that *D*-allo-Ile is used by the former while additional *D*-Leu and *D-*Val are used by the latter fungi. To test whether *D*-Leu and or *D-*Val could be uptaken by *B. bassiana*, feeding assays were conducted with the final concentrations of either *D*-type AA at 100 μg/ ml and 500 μg/ml in SDB, respectively. As a result, no additional peak was detected after fermentations (Fig. S5B), suggesting that the NRPS BesA might have no preference to upload *D-*type AAs at least when *B. bassiana* was grown in SDB.

### Distinct contribution of BVDs to fungal virulence against insect hosts

To determine whether the production of BVDs would contribute to fungal virulence, insect bioassays were first conducted with the WT strains of three fungal species. As a results, in contrast to *B. bassiana* and *B. brongniartii*, *C. militaris* could barely infect and kill fruit flies (*Drosophila melanogaster*) (Fig. 6A). Thus, the WT and PKS gene null mutant of *B. bassiana* and *B. brongniartii* were used for topical infection of both the female adults of fruit fly and the last instar larvae of wax moth (*Galleria mellonella*). The results indicated that, relative to the WT, BVD non-producing mutants of Δ*besB* (Log Rank test, χ^2^=26.01; *P* < 0.0001) and Δ*BrbesB* (χ^2^=36.81; *P* < 0.0001) had significantly reduced virulence against fruit flies (Fig. 6B). However, no statistical difference (*P* > 0.05) was observed between WT and respective mutants against the wax moth larvae (Fig. 6C). The results therefore suggested that the contribution of BVD to fungal virulence might be dependent on the susceptibility of insect species.

**FIG 6.**
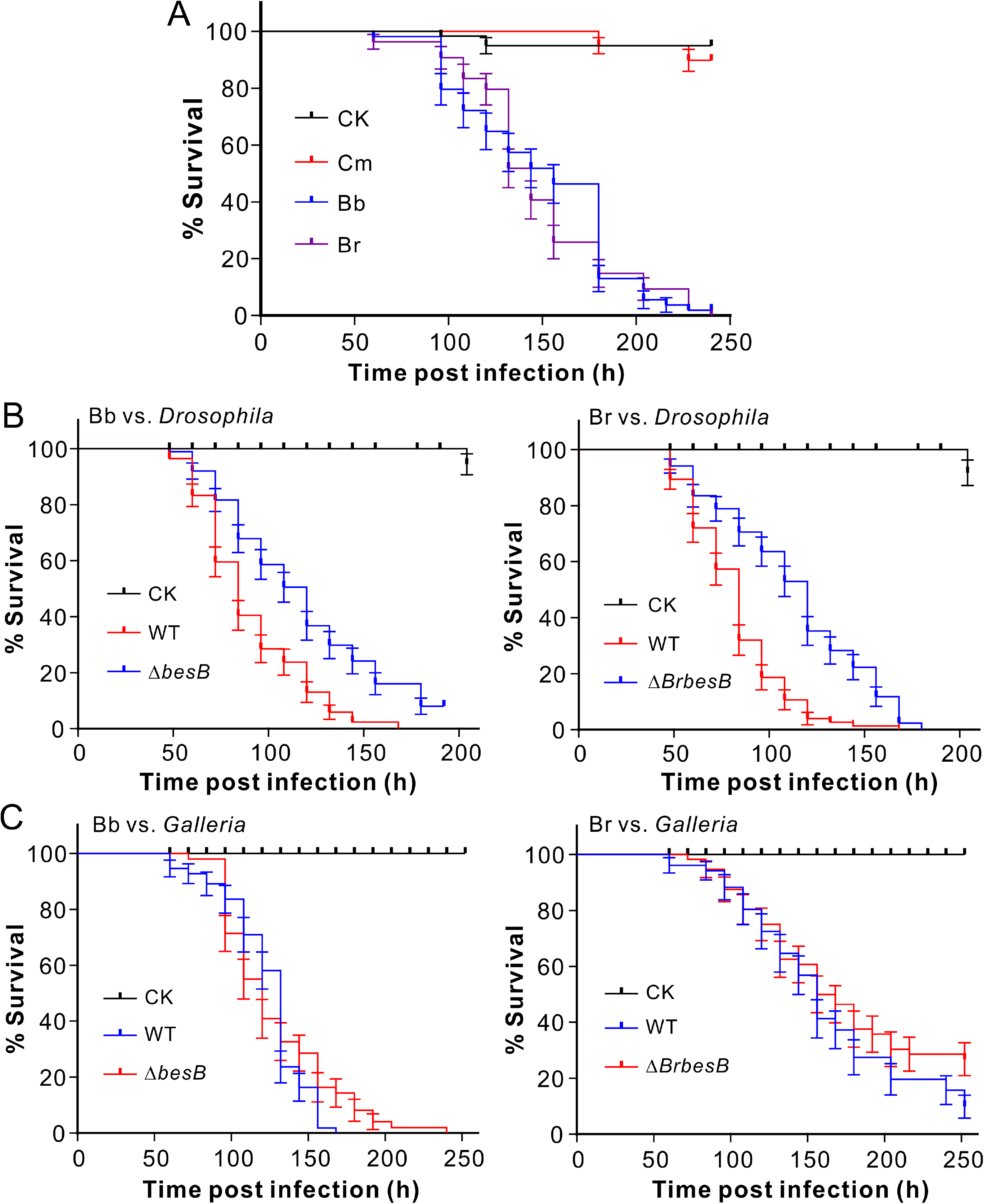
Insect survivals. (A) Survivals of fruit flies after topical infection with the spore suspensions of different WT fungi. (B) Survivals of fruit flies after topical infection with the spore suspensions of the WT and mutant fungi. (C) Survivals of the wax moth larvae after topical infection with the spore suspensions of the WT and mutant fungi. Fungal species are: *C. militaris* (Cm), *B. bassiana* (Bb) and *B. brongniartii* (Br). CK, control.

## DISCUSSION

Fungal secondary metabolite production is largely associated with fungal speciation, the typical feature and basis of fungal chemotaxonomy (27). For example, different structural destruxins are similarly produced by different toxin-producing species of *Metarhizium* fungi (6). The analogous mycotoxin fumonisins can be produced different *Fusarium* species that the early diverged species (e.g., *F. oxysporum* and *F. anthophilum*) produce C-type fumonisins (with a 19-carbon-atom chain) whereas the recently diverged species (e.g., *F. verticillioides* and *F. ujikuroi*) predominantly produce B fumonisins (with a 20-carbon-atom chain) (28). We report in this study that the conserved BGC responsible for BVD biosynthesis produces structurally different analogs in three closely-related Cordycipitaceae fungi. It was found that two *Beauveria* species produce distinct type of BVD analogs whereas the more divergent species *B. brongniartii* and *C. militaris* produce similar compounds. Even the sexual stage of *Beauveria* species has been clarified to be *Cordyceps* spp. (29), different studies have shown that *B. bassiana* is more closely related with *B. brongniartii* than with *C. militaris* (19, 24, 30). The underlying mechanism of BGC functional divergence remains to be determined in different species. This uncommon case would at least question the reliability of fungal chemotaxonomy, especially when using a particular compound as a biomarker (31).

The evolution of fungal BGCs is still intriguing in terms of their disparate distributions in closely related or unrelated fungal species. It has been reported that BVD L and La could be produced by *P. fumosorosea* (syn. *I. fumosorosea*) (8). However, our analysis with the obtained genome information revealed that the BVD BGC is missing in the genomes of *I. fumosorosea* as well as the closely-related species such as *C. confragosa* and *C. cicadae* of Cordycipitaceae fungi (19, 30). Likewise, the conserved BGC for bibenzoquinone oosporein production is present in *B. bassiana* and *B. brongniartii* but absent from *C. militaris* (32). Considering that horizontal gene transfer (HGT) of BGCs frequently occurred among fungal species (1), high conservation between BVD and emericellamide BGCs would suggest the transfer of the cluster from *Aspergillus* to *Cordyceps* fungi along with a terminal truncation of the NRPS gene (i.e. the loss of last two modules) and BGC losses in different species such as *C. confragosa* and *C. cicadae*. An additional support of HGT event from *A. nidulans* to *C. militaris* is the presence of the conserved BGC for the joint production of cordycepin and its protection molecule pentostatin (33).

The structure diversity of BVD analogs is determined by assembly with alternate building blocks of the 3-hydroxy FAs biosynthesized by the PKS enzyme and AAs uploaded and assembled by NRPS synthetase. It has been clarified that each A-domain of NRPS is responsible for the uptake of specific AA residues (1). In contrast to phylogenetic relationships, A-domain signature prediction would support the finding that *B. brongniartii* and *C. militaris* produce compounds with similar AA residues. Even not being detected in this study, a previous report has indicated that *B. bassiana* could also produce beauverolide D, E, Ea, F and Fa that all contain a 3-hydroxy FA of HMOA (26). Taken together with the detection of BVDs Ba, Ca and Ka (all containing a HMDA) in this study, the observations suggest that BesB could produce both HMDA and HMOA like CmBesB and BrBesB. Thus, the PKS gene might not determine the specificity of alternative BVD formations in different fungi. To some extent, it is supported from our phylogenetic analysis of PKS domains (i.e., KS, AT, KR and ER) that may control the reaction of chain elongation, selective tethering or incorporation of intermediates are all congruent with fungal speciation relationships.

The EasB of *A. nidulans* produces 3-hydroxy FAs HDMDA and HTMDA (22), which differs from the counterparts of three insect pathogens in forming HMDA and HMOA. Consistent with our feeding assays that a 3-hydroyx FA longer than 10 carbon atoms could not be used in *C. militaris*, heterologous expression of the BGC genes of *C. militaris* in *A. nidulans* only resulted in the production of BVD I and III (9), which suggested that both HDMDA and HTMDA produced by *Aspergillus* could not be integrated into the assembly line of the *Cordyceps* enzymes. We also found that the interchangeable replacement of the acyl CoA-ligase gene *besD* with *CmbesD* could not alter the production profile of BVDs in *B. bassiana*. The finding implied that both HMDA and HMOA could be similarly catalyzed to CoA-thioesters by homologous CoA-ligases and the gene might not involve in control of the specificity of analogous BVD productions in different fungi. The exact mechanism involved in producing distinct BVD analogs in different fungi remains to be determined.

The acyltransferase activity is responsible for selection and integration of monomeric substrates into the megasynthases of NRPS or PKS (34). We found that, in contrast to the failure of the acyltransferase mutant Δ*easC* to produce emericellamides, the null mutant of *besC* could still produce BVDs even with reduced amounts when compared with the WT. It is possible that a non-clustered acyltransferase gene (e.g., BBA_08697, 22% identity at AA level with BesC) may play the complementary role for transferring the CoA-ligated HMDA or HMOA into the NRPS assembly line, however, which remains to be investigated.

Structurally, BVDs are additionally different from emericellamides that the formers contain a *D-*type AA residue at position 4. *D*-AAs are occasionally proteinaceous. As being found, a *D-*alanine (*D-*Ala) residue is present in the immunosuppressant agent cyclosporins, which biosynthesis requires the function of a clustered *D-*Ala racemase to convert *L-*Ala to *D-*Ala (5). The racemase-like gene is absent in the BGC for BVD biosynthesis. Our genome survey indicated that different pyridoxal phosphate-dependent (PLP) racemases are present in the genomes of Cordycipitaceae fungi. For example, the SimB-like enzyme is present in *B. bassiana* (BBA_07588, 39% identity), *C. militaris* (CCM_08747, 40%) and *B. brongniartii* (BBO_04243, 39%) that all lie outside of the BVD BGC in each fungus. Considering that the position 4 of each BVD contains alternate *D-*AA (Table 1), different PLP-dependent enzymes may be responsible for the provision of *D*-AA residue for the production of diverse BVDs. Our feeding assays with *D*-Val and *D*-Leu that are frequently used by *C. militaris* and *B. brongniartii* could not change the BVD production profiles of *B. bassiana*. The finding suggests that the A3-domain of BesA may determine the substrate and therefore BVD production specificity rather than the cellular shortage of *D-*type amino acids.

Cyclodepsipeptides produced by insect pathogenic fungi such as beauvericin produced by *B. bassiana* and destruxins produced by *Metarhizium* species are non-selectively insecticidal (6, 35). Except for the identified biological/medicinal activities such as being antiatherogenic (15), chemical biology of BVDs is still elusive. Our previously metabolomics analysis indicated that BVD analogs could be detected in insect hemolymph after the infection of *B. bassiana* for 36 hrs (16). In this study, insect bioassays indicated that the BVD non-producing mutants of *B. bassiana* and *B. brongniartii* were apparently impaired during the infection of *Drosophila* but there were no obvious difference in infection of the wax moth larvae when compared with the WT strains. On top of the selective toxicity of BVDs against different insects, the sensitivity of different insects to BVDs might vary. As being reported, injection of *Galleria* with beauveriolide L could not kill insects at a dosage < 30 μg per insect but could induce insect immune responses (17). The exact biological role(s) of BVDs remain to be determine in the future.

In conclusion, we report the production of distinct BVD analogs independent of fungal phylogenetic associations in three Cordycipitaceae fungi. The specificity of analogous BVD production would be mainly determined by the A-domain selectivity of the NRPS enzyme and the biosynthesis of BVDs is non-canonical that may at least require the gene(s) lying outside of the cluster to provide *D-*type amino acids. The results of this study advance the chemodiversity of fungal secondary metabolisms and may facilitate future chemical ecology studies of cyclodepsipeptides.

## Materials and Methods

### Fungal strains and reagents

The WT strains of *B. bassiana* ARSEF 2860 (abbreviated Bb1) and ARSEF 8028 (Bb2), *B. brongniartii* RCEF 3172 (Br), and *C. militaris* Cm01 (Cm) were used in this study for metabolite production and gene deletions. Both the WT and mutants were maintained on potato dextrose agar (PDA; BD Difco). For BVD production, the WT or mutants was inoculated in SDB (BD Difco) in a rotatory shaker at 25 °C and 180 rpm for 10 days.

### Cluster prediction and genetic manipulations

Whole genome analysis of fungal secondary metabolic gene clusters was performed using the program antiSMASH ver. 5.0 (36). A PKS and NRPS gene cluster of three Cordycipitaceae fungi was found to be similar (40-60% similarity; Table S1) to the BGC of emericellamides in *A. nidulans* (22). To determine the functions of the clustered genes, gene deletions were individually performed by *Agrobacterium*-mediated transformation of *B. bassiana* (37). To generate the deletion vectors, the 5’- and 3’-flanking regions of the target gene were amplified using different primer pairs (Table S16). For example, the primers BesA-U1/BesA-U2 were used to amplify the *besA* upstream region, and the primers BesA-L1/BesA-L2 were used to amplify the *besA* downstream region. The PCR fragments were digested with the restriction enzymes, purified and then cloned into the same enzyme-treated binary vector pDHt-SK-Bar (for resistance against glufosinate ammonium) for transformation of the WT strain of *B. bassiana*. In addition, *BrbesA* and *BrbesB* were individually deleted in *B. brongniartii*, and *CmbesA* and *CmbesB* were disrupted in *C. militaris*, respectively. The drug resistance colonies were verified by PCR, and at least two independent null mutants of each gene were obtained by single spore isolation for BVD production profiling. After preliminary analysis, one null mutant of each gene was used for further experiments.

To examine the potential association with substrate specificity, the Δ*besD* mutant of *B. bassiana* was complemented with the *CmbesD* gene from *C. militaris*. Thus, the full-length open reading frame of *CmbesD* was amplified using both the genomic DNA and cDNA of *C. militaris* as templates by fusion PCRs using the ClonExpress II one step cloning kit (Vazyme, China). The gene was made under the control of the constitutive *gpdB* gene (BBA_05480) promoter (38). The obtained cassette was cloned into the plasmid pDHt-SK-Sur-GpdB (with a *sur* gene for conferring sulfonylurea resistance) (39) for transformation of the Δ*besD* mutant of *B. bassiana*.

### Enzyme structure modulation and phylogenetic analysis

Modulation of the core enzymes BesA and BesB of *B. bassana*, and EasA and EasB of *A. nidulans* was performed using the program antiSMASH (36). A-domain signature of NRPSs BesA and EasA was predicted with the algorithm NRPSpredictor2 (40). The A-domain sequences were retrieved from EasA and BesA, and the AT-, ER- and KR-domains extracted from EasB and BesB-like PKSs. To determine the phylogenetic relationship in association with substrate specificity, the domain sequences were aligned with the program MUSCLE (41) and the neighbor-joining trees were generated using the software MEGA X with a Dayhoff model and 1000 bootstrap replicates (42). In addition, the homologs of BesC and BesD were retrieved from different fungi for phylogenetic analysis.

### BVD production and chromatography analysis

The conidia of the WT and different mutants of *B. bassiana*, *B. brongniartii* and *C. militaris* were harvested from the two-week old PDA plates and suspended in 0.05% Tween 20 to a final concentration of 1 × 10^8^ spores/ml. The spore suspensions were inoculated (1 ml each) into the SDB medium (100 ml in each 250 ml flask) and incubated at 25 °C and 180 rpm in a rotatory shaker for 10 days. The mycelia of each sample were then harvested by filtration, washed twice with sterile water and extracted twice with 50 ml methanol for 30 min under sonication. The extracted samples were concentrated under vacuum, and the crude extract was dissolved in 1.5 ml methanol. There were three replicates for each strain.

Considering that the fruiting bodies of *C. militaris* are mainly consumed as health-beneficial mushrooms (43), the fruiting bodies of this fungus was also induced both on the Chinese Tussah silkmoth (*Antheraea pernyi*) pupae and on rice medium (44). After inoculation for 50 days, the fruiting bodies were harvested and freeze dried for methanol extraction. All samples were centrifuged at 10,000 g for 2 min, and the supernatants were further filtered through a 0.45 μm pore-sized filter prior to HPLC analysis using a LC-20AD system (Shimadzu, Japan) equipped with an SPD-20A UV-visible (UV-Vis) detector and a C_18_ reverse-phase column (particle size, 5 μm; 4.6 × 250 mm; CNW Athena C18, China). Sample aliquots (15 μl each) were eluted with the deionized water (solution A) and acetonitrile (solution B, 0~20 min, 15 to 80% acetonitrile; 20~35 min, 80% acetonitrile) at a flow rate of 1 ml/min and monitored at 194 nm. The column temperature was set at 40°C.

### Compound purification and structure analysis

For purification of BVDs from *B. bassiana*, mycelia were harvested from 12 L of SDB broth and extracted with methanol to generate about approximate 10 g of crude extract. The crude sample was first fractionated using an Inertsil ODS (octadecyl silica) column with a gradient elution of deionized water (solution A) and methanol (solution B, 30~100% methanol). BVDs were concentrated in the 80% methanol fraction, which was further purified using a LC-20AD HPLC system with a C_18_ reverse-phase column (particle size, 5 μm; 10 × 250 mm; CNW Athena C18, China). Samples were eluted with the deionized water (solution A) and acetonitrile (solution B, 80% acetonitrile) at a flow rate of 4 ml/min.

For purification of BVDs produced by *B. brongniartii*, mycelia were collected from 2 L of SDB culture broth and extracted with methanol to generate crude extract for purification with a LC-20AD HPLC system. Three fractions that were rich in BVDs were first collected. Beauveriolide V/VI/VII were concentrated in Fraction 1. Beauveriolide I/III were concentrated in Fraction 2. Beauveriolide II/IX/X and Beauverolide La were concentrated in Fraction 3. These three fractions were further purified with the LC-20AD HPLC system to obtain individual BVDs. The purified compounds were individually subject to 1D NMR analysis in pyridine-*d*5 to collect the ^1^H and ^13^C spectra using a Bruker Avance III-500 system equipped with a 5 mm PABBO BB-^1^H/D probe. Standard pulse parameters were used for all NMR experiments. For the new compound beauveriolide X, 2D NMR spectra were also collected to obtain the information of HSQC (heteronuclear singular quantum correlation), HMBC (heteronuclear multiple bond correlation) and COSY (correlation spectroscopy). All spectrum data were processed using the inbuilt program MestReNova (ver. 9.0.1; Metrelab Research, Santiago de Compostella, Spain).

### Substrate feeding assays

To determine the substrate specificity, three 3-hydroxy FAs with chain length variations, i.e., 3-HDDA, 3-HAD and 3-HAD were ordered (Abcam, Cambridge, UK) and used to feed both Δ*besB* of *B. bassiana* and Δ*CmbesB* of *C. militaris*. The cultures were grown in SDB supplemented with each FA at a final concentration of 100 μg/ml five days post inoculation, and kept for cultivation for additional five days. After incubation, mycelial samples were harvested for metabolite extraction with methanol as described above. Relative to the Δ*CmbesB* sample, additional peak(s) obtained in individual FA feeding experiment were collected and subject to LC-MS analysis. Compound structures were deduced based on the obtained mass data. To determine the effect of *D-*type AA additions on BVD production, *D-*Leu and *D*-Val were used to feed the WT *B. bassiana* strain ARSEF 2860. The spores were inoculated into 250 ml flasks containing 100 ml SDB for four days, and *D-*Leu and *D*-Val were individually added at the final concentrations of 100 μg/ml and 500 μg/ml, respectively. The samples were incubated for another four days and mycelia were harvested and extracted with methanol for HPLC analysis.

### Insect bioassays

To determine if any contribution of BVDs to fungal virulence against insects, the WT strains of *B. bassiana*, *B. brongniartii* and *C. militaris* were first examined using the female adults of *Drosophila melanogaster* by topical infection with spore suspensions (1 × 10^7^ conidia/ml) (45). After verification that *C. militaris* was non-virulent to fruit flies, the WT strains of *B. bassiana* and *B. brongniartii*, and *besB* and *BrbesB* deletion mutants were then assayed in parallel against the fruit flies and the last instar larvae of the wax moth *G. mellonella* as being described before (37). Insect mortality was recorded every 12 hrs and the survival dynamics were compared between WT and mutant of each species by Kaplan–Meier analysis.

## SUPPLEMENTAL MATERIAL

**Table S1** Gene cluster contents in three insect pathogens.

**Table S2** NMR data (^13^ C, 125 MHz; ^1^ H, 500 MHz) of Beauverolide Ba in pyridine-*d*5.

**Table S3** NMR data (^13^ C, 125 MHz; ^1^ H, 500 MHz) of Beauverolide Ca in pyridine-*d*5.

**Table S4** NMR data (^13^ C, 125 MHz; ^1^ H, 500 MHz) of Beauverolide Ka in pyridine-*d*5.

**Table S5** NMR data (^13^ C, 125 MHz; ^1^ H, 500 MHz) of Beauverolide La in pyridine-*d*5.

**Table S6** NMR data (^13^ C, 125 MHz; ^1^ H, 500 MHz) of Beauveriolide I in pyridine-*d*5.

**Table S7** NMR data (^13^ C, 125 MHz; ^1^ H, 500 MHz) of Beauveriolide II in pyridine-*d*5.

**Table S8** NMR data (^13^ C, 125 MHz; ^1^ H, 500 MHz) of Beauveriolide III in pyridine-*d*5.

**Table S9** NMR data (^13^ C, 125 MHz; ^1^ H, 500 MHz) of Beauveriolide V in pyridine-*d*5.

**Table S10** NMR data (^13^ C, 125 MHz; ^1^ H, 500 MHz) of Beauveriolide VI in pyridine-*d*5.

**Table S11** NMR data (^13^ C, 125 MHz; ^1^ H, 500 MHz) of Beauveriolide VII in pyridine-*d*5.

**Table S12** NMR data (^1^ H, 500 MHz) of Beauveriolide IX in pyridine-*d*5.

**Table S13** NMR data (^13^ C, 125 MHz; ^1^ H, 500 MHz) of Beauveriolide X in pyridine-*d*5.

**Table S14** Structure composition of bvd analogs identified in this study.

**Table S15** Putative compositions of the compounds obtained from the mutant Δ*CmbesB* fed with 3-HDA and 3-HOA.

**Table S16** PCR Primers used in this study.

**FIG S1** The structures of beauveriolides (**I**-**X**) and beauverolides (**Ba**, **Ca**, **Ka** and **La**) identified in this study from different *Cordyceps* fungi. HMDA, 3-hydroxy-4-methyldecanoic acid; HMOA, 3-hydroxy-4-methyloctanoic acid.

**FIG S2** 1D and 2D NMR analysis of the compound beauveriolide **X**. NMR data (^13^C, 125 MHz; ^1^H, 500 MHz) were collected in pyridine-*d*5. HSQC, heteronuclear singular quantum correlation. HMBC, heteronuclear multiple bond correlation. COSY, correlation spectroscopy.

**FIG S3** Fruiting-body (FB) induction and metabolite profiling of *C. militaris*. (A) Phenotype of fruiting bodies formed on caterpillar pupa (left) and rice medium. The silkmoth pupae were injected with spore suspensions or the rice medium were inoculated for 50 days. (B) HPLC profiling of metabolite production or non-production by different samples of *C. militaris*. SDB represents the extraction from the mycelial sample harvested from SDB.

**FIG S4** Phylogenetic analysis of the PKS domains and tailoring enzymes involved in bvd biosynthesis in different fungi. Different domains: KS, ketosynthase; AT, acyl transferase; KR, ketoreductase; ER, enoyl reductase. The domains of other PKS or other tailoring enzymes were included to root the tree: OpS1 (BBA_08179), PKS for oosporein biosynthesis in *B. bassiana*; AfoG (AN1026), PKS for Asperfuranone biosynthesis in *A. nidulans*; TRI101 (AAD19745) trichothecene 3-O-acetyltransferase of *Fusarium sporotrichioides*; InpC (AN3490) Acyl-CoA ligase for the biosynthesis of Fellutamide B in *A. nidulans*.

**FIG S5** Substrate feeding assays of *B. bassiana*. (A) HPLC profiling of the PKS gene *besB* deletion mutant after feeding with different demethyl-hydroxy-FAs. Each FA was added at a final concentration of 100 μg/ml. FAs: 3-HDDA, 3-hydroxydodecanoic acid; 3-HAD, 3-Hydroxydecanoic acid; 3-HOA, 3-hydroxyoctanoic acid. (B) HPLC profiling of *B. bassiana* after feeding with different concentrations of *D*-Val and *D-*Leu. The spores of the *B. bassiana* WT strain ARSEF 2860 were inoculated in SDB for four days and then the *D*-type amino acids were added at the final concentrations as indicated for another four days. Mock control was incubated in SDB without the addition of any *D*-type amino acid.

## ACKNOWLEDGMENTS

We thank Mr. Shizhen Bu for his help with NMR analysis of compound structures. This work was supported by the National Natural Science Foundation of China (grant 31530001), Chinese Academy of Sciences (grant QYZDJ-SSW-SMC028), and the Shanghai Academic/Technology Research Leader Program (grant 18XD1404500).

## Reference

1. Keller NP. 2019. Fungal secondary metabolism: regulation, function and drug discovery. Nat Rev Microbiol 17:167–180.

2. Sivanathan S, Scherkenbeck J. 2014. Cyclodepsipeptides: a rich source of biologically active compounds for drug research. Molecules 19:12368–12420.

3. Survase SA, Kagliwal LD, Annapure US, Singhal RS. 2011. Cyclosporin A--a review on fermentative production, downstream processing and pharmacological applications. Biotechnol Adv 29:418–435.

4. Liu BL, Tzeng YM. 2012. Development and applications of destruxins: a review. Biotechnol Adv 30:1242–1254.

5. Yang X, Feng P, Yin Y, Bushley K, Spatafora JW, Wang C. 2018. Cyclosporine biosynthesis in *Tolypocladium inflatum* benefits fungal adaptation to the environment. mBio 9:e01211–01218.

6. Wang B, Kang Q, Lu Y, Bai L, Wang C. 2012. Unveiling the biosynthetic puzzle of destruxins in *Metarhizium* species. Proc Natl Acad Sci USA 109:1287–1292.

7. Elsworth JF, Grove JF. 1977. Cyclodepsipeptides from *Beauveria bassiana* Bals. Part 1. Beauverolides H and I. J Chem Soc, Perkin Trans 1:270–273.

8. Jegorov A, Sedmera P, Matha V, Simek P, Zahradnickova H, Landa Z, Eyal J. 1994. Beauverolides L and La from *Beauveria tenella* and *Paecilomyces fumosoroseus*. Phytochemistry 37:1301–1303.

9. Wang X, Gao YL, Zhang ML, Zhang HD, Huang JZ, Li L. 2020. Genome mining and biosynthesis of the Acyl-CoA:cholesterol acyltransferase inhibitor beauveriolide I and III in *Cordyceps militaris*. J Biotechnol 309:85–91.

10. Jegorov A, Paizs B, Kuzma M, Zabka M, Landa Z, Sulc M, Barrow MP, Havlicek V. 2004. Extraribosomal cyclic tetradepsipeptides beauverolides: profiling and modeling the fragmentation pathways. J Mass Spectrom 39:949–960.

11. Matsuda D, Namatame I, Tomoda H, Kobayashi S, Zocher R, Kleinkauf H, Omura S. 2004. New beauveriolides produced by amino acid-supplemented fermentation of *Beauveria* sp. FO-6979. J Antibiot (Tokyo) 57:1–9.

12. Nakaya S, Mizuno S, Ishigami H, Yamakawa Y, Kawagishi H, Ushimaru T. 2012. New rapid screening method for anti-aging compounds using budding yeast and identification of beauveriolide I as a potent active compound. Biosci Biotechnol Biochem 76:1226–1228.

13. Witter DP, Chen Y, Rogel JK, Boldt GE, Wentworth P, Jr. 2009. The natural products beauveriolide I and III: a new class of beta-amyloid-lowering compounds. ChemBioChem 10:1344–1347.

14. Ohshiro T, Kobayashi K, Ohba M, Matsuda D, Rudel LL, Takahashi T, Doi T, Tomoda H. 2017. Selective inhibition of sterol O-acyltransferase 1 isozyme by beauveriolide III in intact cells. Sci Rep 7:4163.

15. Namatame I, Tomoda H, Ishibashi S, Omura S. 2004. Antiatherogenic activity of fungal beauveriolides, inhibitors of lipid droplet accumulation in macrophages. Proc Natl Acad Sci U S A 101:737–742.

16. Xu Y-J, Luo F, Gao Q, Shang Y, Wang C. 2015. Metabolomics reveals insect metabolic responses associated with fungal infection. Anal Bioanal Chem 407:4815–4821.

17. Vilcinskas A, Jegorov A, Landa Z, Gotz P, Matha V. 1999. Effects of beauverolide L and cyclosporin A on humoral and cellular immune response of the greater wax moth, *Galleria mellonella*. Comp Biochem Physiol C Pharmacol Toxicol Endocrinol 122:83–92.

18. Nagai K, Doi T, Sekiguchi T, Namatame I, Sunazuka T, Tomoda H, Omura S, Takahashi T. 2006. Synthesis and biological evaluation of a beauveriolide analogue library. J Comb Chem 8:103–109.

19. Shang YF, Xiao GH, Zheng P, Cen K, Zhan S, Wang CS. 2016. Divergent and convergent evolution of fungal pathogenicity. Genome Biol Evol 8:1374–1387.

20. Zheng P, Xia Y, Xiao G, Xiong C, Hu X, Zhang S, Zheng H, Huang Y, Zhou Y, Wang S, Zhao GP, Liu X, St Leger RJ, Wang C. 2011. Genome sequence of the insect pathogenic fungus *Cordyceps militaris*, a valued traditional Chinese medicine. Genome Biol 12:R116.

21. Xiao G, Ying SH, Zheng P, Wang ZL, Zhang S, Xie XQ, Shang Y, St Leger RJ, Zhao GP, Wang C, Feng MG. 2012. Genomic perspectives on the evolution of fungal entomopathogenicity in *Beauveria bassiana*. Sci Rep 2:483.

22. Chiang YM, Szewczyk E, Nayak T, Davidson AD, Sanchez JF, Lo HC, Ho WY, Simityan H, Kuo E, Praseuth A, Watanabe K, Oakley BR, Wang CC. 2008. Molecular genetic mining of the *Aspergillus* secondary metabolome: discovery of the emericellamide biosynthetic pathway. Chem Biol 15:527–532.

23. Wang C, Wang S. 2017. Insect pathogenic fungi: genomics, molecular interactions, and genetic improvements. Annu Rev Entomol 62:73–90.

24. Kepler RM, Luangsa-Ard JJ, Hywel-Jones NL, Quandt CA, Sung GH, Rehner SA, Aime MC, Henkel TW, Sanjuan T, Zare R, Chen M, Li Z, Rossman AY, Spatafora JW, Shrestha B. 2017. A phylogenetically-based nomenclature for Cordycipitaceae (Hypocreales). IMA Fungus 8:335–353.

25. Mei LJ, Chen M, Shang Y, Tang G, Tao Y, Zeng L, Huang B, Li Z, Zhan S, Wang CS. 2020. Population genomics and evolution of a fungal pathogen after releasing exotic strains to control insect pests for 20 years. ISME J 14:1422–1434.

26. Elsworth JF, Grove JF. 1980. Cyclodepsipeptides from *Beauveria bassiana*. Part 2. Beauverolides A to F and their relationship to isarolide. J Chem Soc, Perkin Trans 1 1:1795–1799.

27. Frisvad JC, Andersen B, Thrane U. 2008. The use of secondary metabolite profiling in chemotaxonomy of filamentous fungi. Mycol Res 112:231–240.

28. Proctor RH, Van Hove F, Susca A, Stea G, Busman M, van der Lee T, Waalwijk C, Moretti A, Ward TJ. 2013. Birth, death and horizontal transfer of the fumonisin biosynthetic gene cluster during the evolutionary diversification of *Fusarium*. Mol Microbiol 90:290–306.

29. Li ZZ, Li CR, Huang B, Fan MZ. 2001. Discovery and demonstration of the teleomorph of *Beauveria bassiana* (Bals.) Vuill., an important entomogenous fungus. Chinese Sci Bull 46:751–753.

30. Lu Y, Luo F, Cen K, Xiao G, Yin Y, Li C, Li Z, Zhan S, Zhang H, Wang C. 2017. Omics data reveal the unusual asexual-fruiting nature and secondary metabolic potentials of the medicinal fungus *Cordyceps cicadae*. BMC Genomics 18:668.

31. Isaka M, Sappan M, Jennifer Luangsa-Ard J, Hywel-Jones NL, Mongkolsamrit S, Chunhametha S. 2011. Chemical taxonomy of *Torrubiella* s. lat.: zeorin as a marker of Conoideocrella. Fungal Biol 115:401–405.

32. Feng P, Shang Y, Cen K, Wang C. 2015. Fungal biosynthesis of the bibenzoquinone oosporein to evade insect immunity. Proc Natl Acad Sci USA 112:11365–11370.

33. Xia YL, Luo FF, Shang YF, Chen PL, Lu YZ, Wang CS. 2017. Fungal cordycepin biosynthesis is coupled with the production of the safeguard molecule pentostatin. Cell Chem Biol 24:1479–1489.

34. Dunn BJ, Khosla C. 2013. Engineering the acyltransferase substrate specificity of assembly line polyketide synthases. J R Soc Interface 10:20130297.

35. Xu Y, Orozco R, Wijeratne EM, Gunatilaka AA, Stock SP, Molnar I. 2008. Biosynthesis of the cyclooligomer depsipeptide beauvericin, a virulence factor of the entomopathogenic fungus *Beauveria bassiana*. Chem Biol 15:898–907.

36. Blin K, Shaw S, Steinke K, Villebro R, Ziemert N, Lee SY, Medema MH, Weber T. 2019. antiSMASH 5.0: updates to the secondary metabolite genome mining pipeline. Nucleic Acids Res 47:W81–w87.

37. Cen K, Li B, Lu YZ, Zhang SW, Wang CS. 2017. Divergent LysM effectors contribute to the virulence of *Beauveria bassiana* by evasion of insect immune defenses. PLoS Pathog 13:e1006604.

38. Liao XG, Fang WG, Zhang YJ, Fan YH, Wu XW, Zhou Q, Pei Y. 2008. Characterization of a highly active promoter, PBbgpd, in *Beauveria bassiana*. Curr Microbiol 57:121–126.

39. Zhang S, Fan Y, Xia YX, Keyhani NO. 2010. Sulfonylurea resistance as a new selectable marker for the entomopathogenic fungus *Beauveria bassiana*. Appl Microbiol Biotechnol 87:1151–1156.

40. Röttig M, Medema MH, Blin K, Weber T, Rausch C, Kohlbacher O. 2011. NRPSpredictor2—a web server for predicting NRPS adenylation domain specificity. Nucleic Acids Res 39:W362–W367.

41. Edgar RC. 2004. MUSCLE: multiple sequence alignment with high accuracy and high throughput. Nucleic Acids Res 32:1792–1797.

42. Kumar S, Stecher G, Li M, Knyaz C, Tamura K. 2018. MEGA X: Molecular Evolutionary Genetics Analysis across Computing Platforms. Mol Biol Evol 35:1547–1549.

43. Chen B, Sun YL, Luo FF, Wang CS. 2020. Bioactive metabolites and potential mycotoxins produced by *Cordyceps* fungi: A review of safety. Toxins (Basel) 12:410.

44. Xiong CH, Xia YL, Zheng P, Wang CS. 2013. Increasing oxidative stress tolerance and subculturing stability of *Cordyceps militaris* by overexpression of a glutathione peroxidase gene. Appl Microbiol Biotechnol 97:2009–2015.

45. Huang A, Lu M, Ling E, Li P, Wang CS. 2020. A M35 family metalloprotease is required for fungal virulence against insects by inactivating host prophenoloxidases and beyond. Virulence 11:222–237.

46. Isogai A, Kanaoka M, Matsuda H, Hori Y, Suzuki A. 1978. Structure of a new cyclodepsipeptide, beauverilide A from *Beauveria bassiana*. Agricultural and Biological Chemistry 42:1797–1798.

47. Grove JF. 1980. Cyclodepsipeptides from *Beauveria bassiana*. Part 3. The isolation of beauverolides Ba, Ca, Ja, and Ka. J Chem Soc, Perkin Trans 1:2878–2880.

48. Kuzma M, Jegorov A, Kacer P, Havlicek V. 2001. Sequencing of new beauverolides by high-performance liquid chromatography and mass spectrometry. J Mass Spectrom 36:1108–1115.

49. Jegorov A, Hajduch M, Sulc M, Havlicek V. 2006. Nonribosomal cyclic peptides: specific markers of fungal infections. J Mass Spectrom 41:563–576.

50. Mochizuki K, Ohmori K, Tamura H, Shizuri Y, Nishiyama S, Miyoshi E, Yamamura S. 1993. The structures of bioactive cyclodepsipeptides, beauveriolides I and II, metabolites of entomopathogenic fungi *Beauveria* sp. Bull Chem Soc Jpn 66:3041–3046.

51. Namatame I, Tomoda H, Tabata N, Si S, Omura S. 1999. Structure elucidation of fungal beauveriolide III, a novel inhibitor of lipid droplet formation in mouse macrophages. J Antibiot (Tokyo) 52:7–12.

